# High-Resolution Structures of Tobacco Mosaic Virus Disks from Cryo-Electron Microscopy

**DOI:** 10.1101/2025.01.15.633274

**Authors:** Ismael Abu-Baker, Artur P. Biela, Sachin N. Shah, Jonathan G. Heddle, George P. Lomonossoff, Amy Szuchmacher Blum

**Affiliations:** Department of Chemistry, McGill University, Montréal, Québec, Canada; Jagiellonian University, SOLARIS National Synchrotron Radiation Centre, Krakow, Poland; Jagiellonian University, Malopolska Centre of Biotechnology, Krakow, Poland; Department of Biochemistry and Metabolism, John Innes Centre, Norwich Research Park, Norwich, UK; School of Biological and Biomedical Sciences, Durham University, Durham, UK

## Abstract

Tobacco mosaic virus has been involved in many important developments in virology, structural biology, and biotechnology. Despite decades of study, several key aspects of the viral assembly mechanism remain unclear, particularly the structure of the coat protein disk that initiates virus assembly by interacting with the viral RNA. Here we report the first cryo-electron microscopy structures of the coat protein disk under conditions typically used for *in vitro* virus assembly. We identify 1-, 2-, and 3-layered disks, all of which differ significantly from previous structures obtained by X-ray crystallography. These new models lead to a revised viral assembly mechanism. We also compare coat proteins produced in bacteria and plants to better understand the effect of N-terminal acetylation.

## Introduction

As the first virus discovered, tobacco mosaic virus (TMV) has played an important role in the development of virology, structural biology, and biotechnology.^1,2^ TMV was one of the first samples studied in the early days of X-ray fibre diffraction and electron microscopy, and it has since become a standard sample for validating new instruments and characterization methods.^3–5^ TMV has also been extensively investigated for biotechnological applications such as templated metal nanomaterials, targeted drug delivery, and artificial light-harvesting arrays.^6–8^

TMV is a helical rod-shaped virus consisting of a single RNA strand protected by numerous copies of the coat protein. Under alkaline conditions and low ionic strength, TMV coat protein (TMVP) exists as a mixture of monomers and small oligomers known as “A-protein”. Non-helical, 2-layer disks with 17 monomers per layer form near neutral pH. At high ionic strength these disks form non-helical stacks. Below pH 6.5, disks begin to self-assemble into helical rods that closely resemble the intact virus but lack RNA.^9,10^ The transition from A-protein to disks and then to helical rods is regulated by protonation of clusters of carboxylate and carboxamide side chains called “Caspar carboxylates”.^11^ Protonation of these clusters reduces electrostatic repulsion, allowing further assembly. The exact structure of the 2-layer disk in solution and how it transitions to the helical rod remains unclear. Based on the crystal structure of a 4-layer disk aggregate, two different 2-layer disks have been proposed: the AA-disk, which has *C*_*2*_-symmetry, and the polar AB-disk, which does not have *C*_*2*_-symmetry (Fig. 1).^12,13^ TMVP features a bundle of four α-helices, denoted as left/right radial (LR/RR) and left/right slewed (LS/RS) based on their relative orientations within the helical rod. The alpha helices in the AA-disk are oriented in the plane of the disk, while those in the AB-disk are tilted out of the disk plane. In the present study we use head/tail notation to describe interactions between disk layers in a more generalizable manner. As shown in Fig. 1D, the head of the native TMV rod is set as the 3′-end and the tail is the 5′-end.^14^ This defines the surface of the monomer with the radial helices as the tail and the surface with slewed helices as the head. The AA-disk therefore becomes a head-head disk, referring to the interface between the two layers, and the AB-disk becomes a head-tail disk. The 4-layer aggregate was crystallized under conditions (0.3 M (NH_4_)_2_SO_4_, 0.1 M Tris/HCl, pH 8.0) that are quite different from typical *in vitro* assembly conditions, which raises concerns about its relevance to virus assembly.^13^

**Figure 1.**
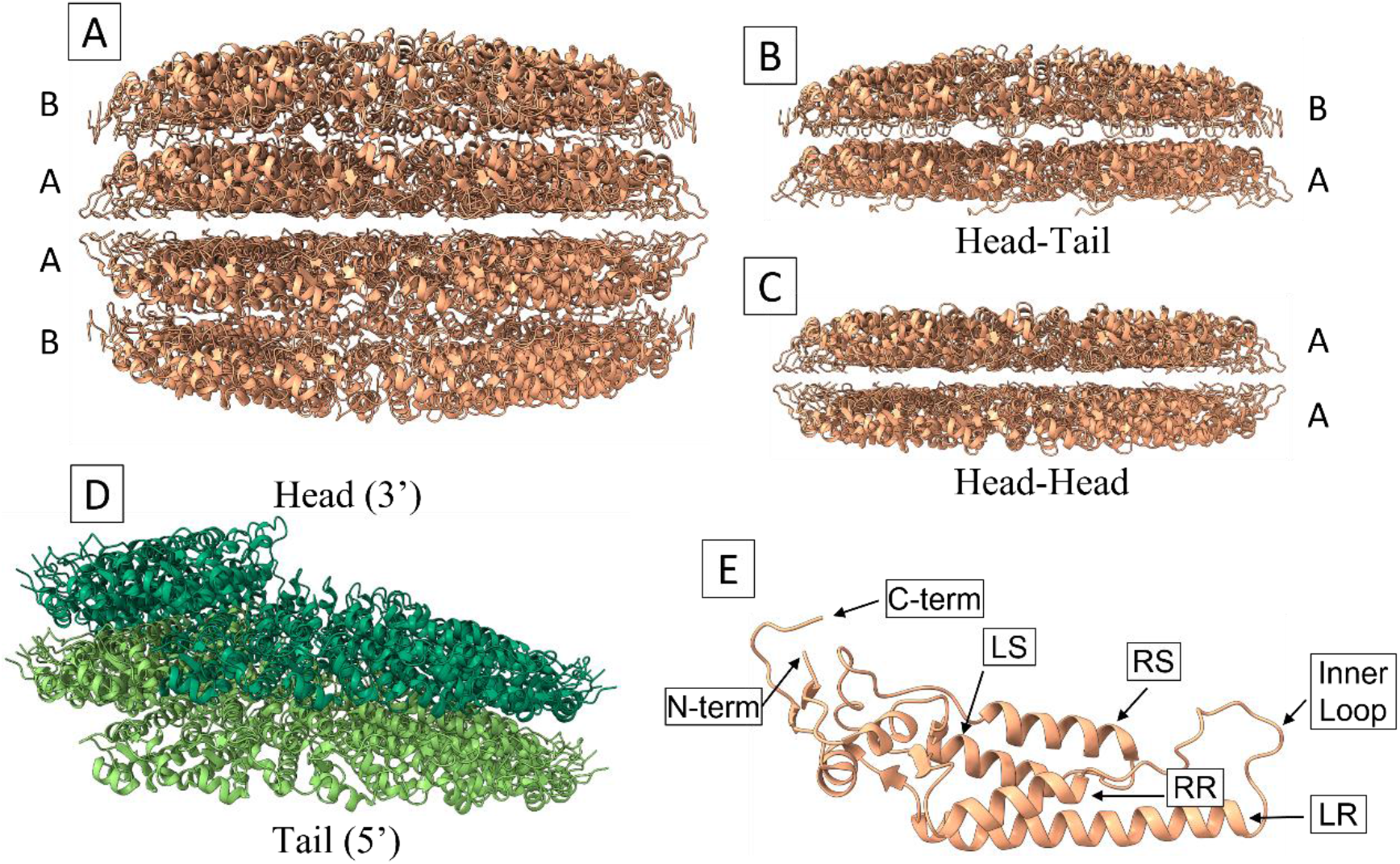
TMV models from literature. (A) 4-layer disk aggregate from PDB 1EI7. (B) AB-disk. (C) AA-disk. (D) Two turns of the helical rod model (PDB 6SAE) showing how the head and tail notation is defined. (E) TMVP monomer with key regions labeled: termini, helices, and inner loop.

While early studies of stacked disks suggested that the disks were polar, more recent studies with higher resolution data as well as antibody binding experiments provide strong evidence that the 2-layer disks in stacked disk assemblies have dihedral symmetry, as in the AA-disk of the 4-layer aggregate (Fig. 1).^15–17^ It has been proposed that the polar 2-layer disk transitions to a metastable 2-turn helix, called a “proto-helix”, which then proceeds to the full helical rod.^18^ There is limited experimental evidence for the existence of the proto-helix or AB-disk as stable species in solution, with only one early cryo-EM study identifying the proto-helix and polar disk in solution within a narrow pH range.^19^ While some of the particles imaged in the study are consistent with the 2D projections of proto-helix and AB-disk models, the resolution and image analysis methods available at the time were far below today’s standards. Additionally, the previous cryo-EM study did not lead to protein models, so there is limited structural insight that can be gained from it.

Considering that almost all structural information for TMVP disks comes from X-ray crystallography and sedimentation experiments, there is a clear need to study disks in solution as close as possible to typical virus assembly conditions. It has also been established that TMVP produced in plants and yeast efficiently encapsidates viral RNA *in vitro* under a range of assembly conditions, while protein produced in bacteria generally fails to encapsidate RNA *in vitro*.^20,21^ This difference in assembly behaviour is attributed to N-terminal acetylation which occurs in plants and some yeast, but not in bacteria. N-terminal acetylation is known to change protein stability and folding in other proteins.^22^ However, the precise structural changes caused by the N-terminal modification in TMVP and how any such structural changes relate to RNA encapsidation have not been determined. In this study, we perform cryo-EM on wild-type TMVP disks from proteins produced in both bacteria and plants with the goal of elucidating the behaviour of TMVP disks in solution and investigating the differences in assembly behaviour of plant-(pTMVP) and bacterially-derived (bTMVP) protein. The cryo-EM results confirm, for the first time, the existence of 1- and 3-layer pTMVP disks, providing new insight into the transition from disks to helical rod.

## Results and Discussion

Cryo-EM data was collected for WT-TMVP produced in both plants and *E. coli*. All the final structures had approximately 3 Å resolution. Samples were stored at pH 5.0 and prepared by dialysis against 75 mM sodium potassium phosphate buffer at pH 7.2 to replicate conditions commonly used for *in vitro* assembly of TMVP with RNA.

### bTMVP Stacked Disks

Bacterially-produced TMVP showed primarily stacked disks with *C*_*2*_-symmetry (Fig. 2A). This behaviour matches previous reports that bTMVP forms stacked disks under acidic conditions that do not disassemble upon dialysis to alkaline pH.^23^ It has been suggested that stacked disks observed in TEM of pTMVP are due to drying effects, but this is not the case in bTMVP.^24^ The disks are head-to-head assemblies, closely resembling the AA-disk. The outer and middle radius of the disk matches very well with the A-disk from the literature. Residues 92-108 of the flexible inner loop were unresolved in the cryo-EM map. Although side chain conformations are mostly conserved, the backbone of the LR helix differs towards the inner radius, starting near Asp116. R112 and R113 show significantly different side chain conformations compared to the literature model. At the start of the inner loop, just after the RR helix, Thr89 and Arg90 were resolved and differ from the AA-disk in backbone and side chain position.

**Figure 2.**
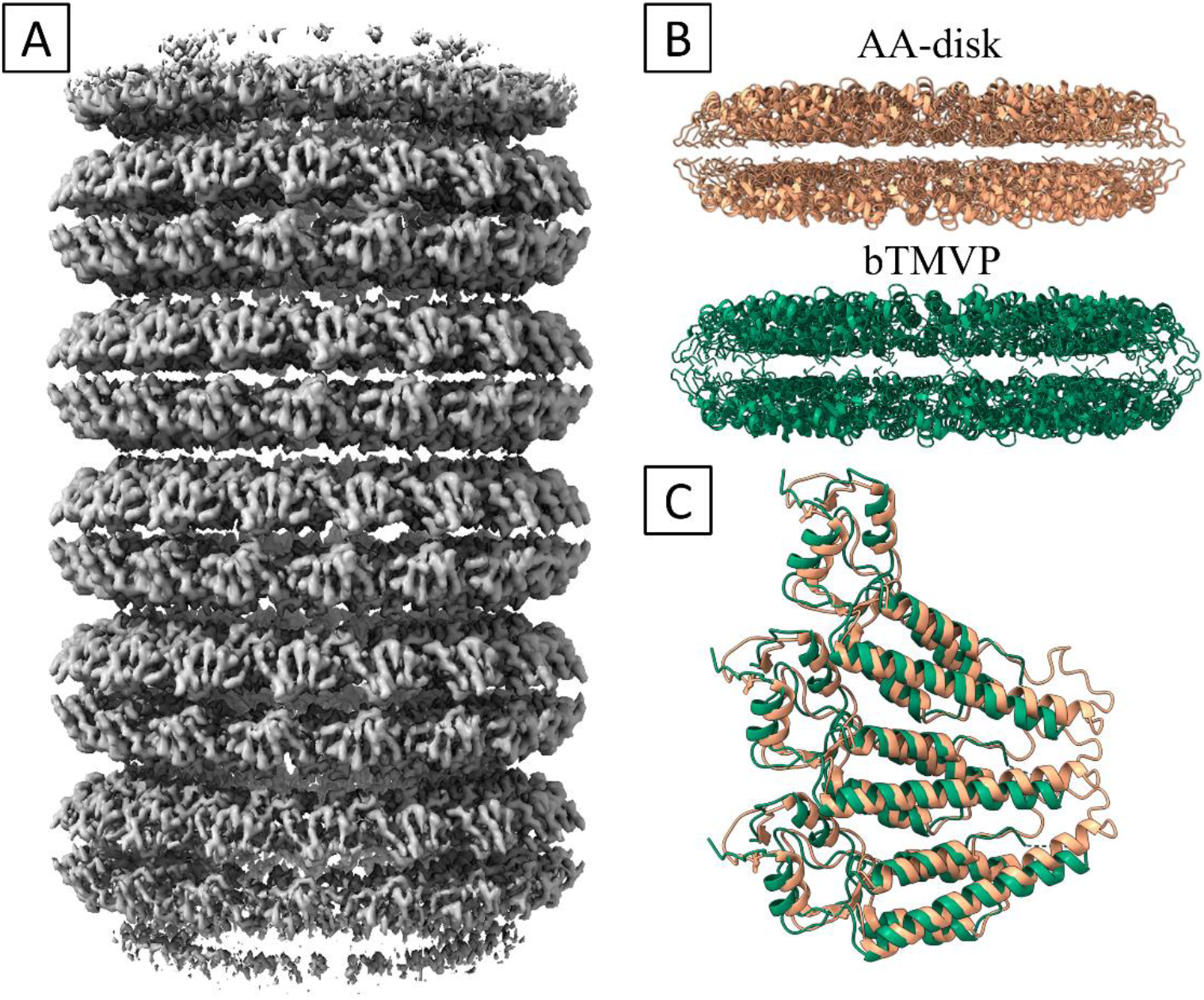
(A) cryo-EM map of bTMVP stacked disks. (B) Side views of AA-disk and refined bTMVP model. (C) Face view of trimer from aligned AA- and bTMVP disks.

Two clusters of protein-protein contacts (within 4 Å of each other) were identified at the head-head interface holding the bilayer disk together. Gln57-Gln57′ and Val60/Thr59-Thr59′/Val60′ form contact pairs at the loop near the outer radius. Near the middle radius, Arg46 interacts with Gln39′′ and Val43′′ on a second opposing subunit (Fig. 3A, B). Axial contacts were also identified at the tail-tail interface, contributing to the stacking of bilayer disks. Arg134 at the outer radius of the LR-helix is oriented along the stacked disk axis on the outer face of each bilayer disk and interacts with Arg134′ on the outer face of the neighbouring disk. This Arg-Arg interaction is stabilized by a Glu131 carboxylate adjacent to each Arg134 side chain. Slightly inward of Arg134, Val130-Val130′ pairs are also within 4 Å of each other. At the same radius on the adjacent RR-helix, a hydrophobic contact is apparent between Ala74/Val75 and Val75′/Ala74′. While not directly forming axial contacts themselves, Thr134 and Glu131 form a lateral bridge between the Arg134 and Val75/Ala74 axial contacts. Towards the inner radius, Arg112 residues appear to engage in similar Arg-Arg interactions, although the side chains were not as well resolved (Fig. 3C-E). Like the Arg134 pairs, Arg112 interactions are stabilized by proximal Asp116 carboxylates. These Arg-Arg interactions have not been previously reported for stacked disks, but Arg-Arg pairs and clusters are well-documented in the literature. Solvent-exposed Arg-Arg pairs can form favourable contacts through pi-pi interactions despite their like-charges, especially in the presence of suitable counter-ions.^25–28^ The N-terminal amine only appears to have contacts with Tyr152 and the backbone oxygen of Thr153. The last three residues of the C-terminus were unresolved, so we cannot conclusively determine if an interaction occurs between the oppositely charged termini, although the flexibility indicates that it is unlikely to be a strong interaction if it does occur. Lateral contacts within each layer are primarily between hydrophobic/amphiphilic residues towards the outer radius and hydrogen bonding or salt bridge interactions towards the inner radius. Further data collection is underway to attain reconstructions of non-stacked disks by avoiding storage at low pH. Preliminary data indicates that bTMVP exists as a mixture of 1-, 2-, 3-, and 4-layer disks if it is not exposed to acidic conditions (Fig. S1).

**Figure 3.**
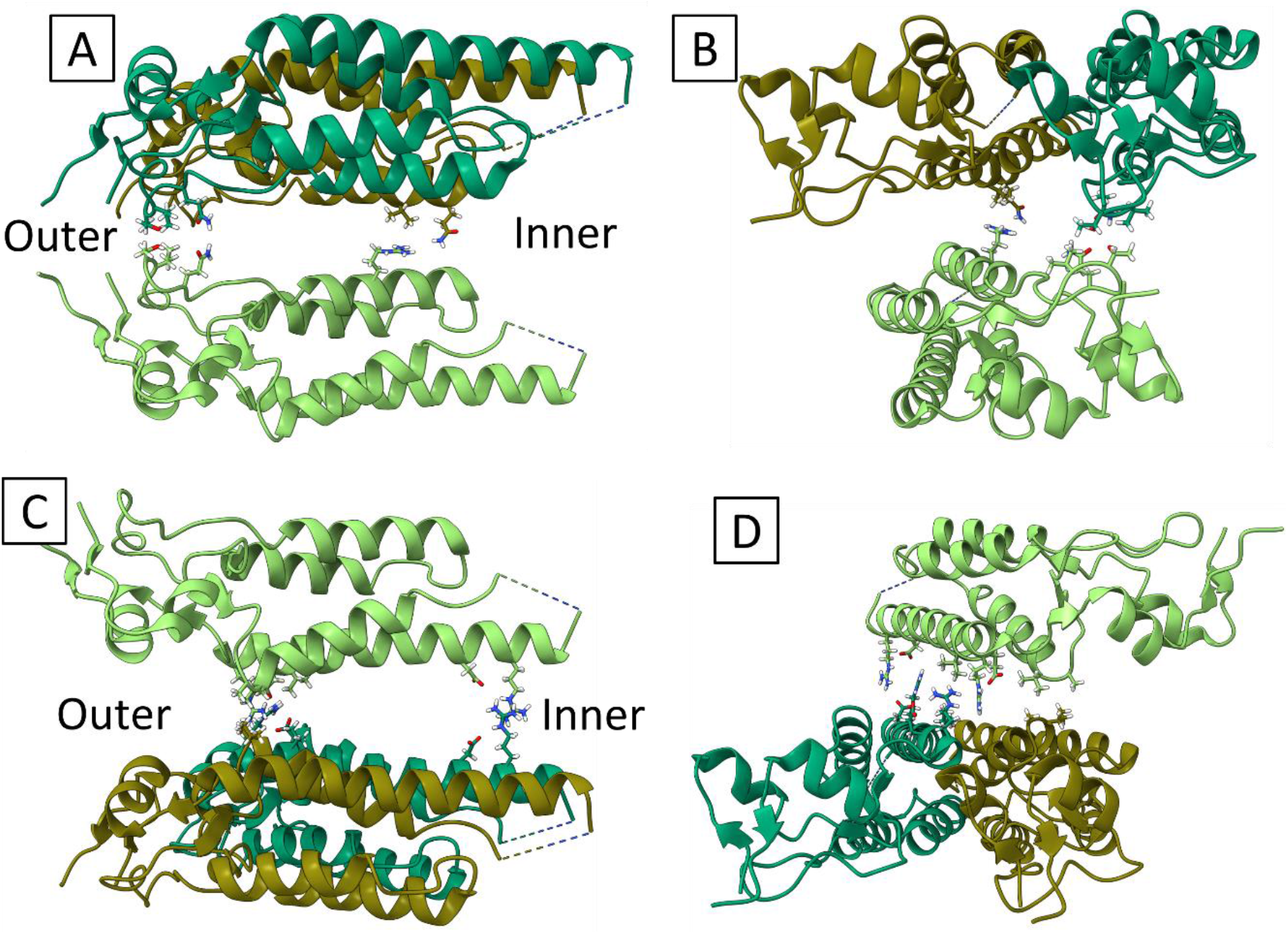
Protein-protein axial contacts of the bTMVP stacked disks. (A) Side view of head-head interface. (B) Radial view of head-head interface. (C) Side view of tail-tail interface. (D) Radial view of tail-tail interface. Each monomer is coloured a different shade of green.

### Plant-derived TMVP

Plant-derived tobacco mosaic virus coat protein formed three distinct non-helical disk species. 50.9% of the particles were 1-layer disks, 17.4% were 2-layer disks, and 31.6% were 3-layer disks (Fig. 4). This is the first time a 1-layer wild-type TMVP disk has been reported, although 1-layer and 3-layer disks have been observed in related viruses, such as barley stripe mosaic virus.^29^ A 3-layer disk or 3-turn helix has been theorized to be the stable nucleating species of the helical rod since the 1960’s.^30^ Evidence for a 3-layer species has mostly been inferred from sedimentation experiments. Recently, peaks matching the masses of 1- and 3-layer disks were observed in a mass spectrometry study of TMVP disks by Bischoff *et al*. and negative stain TEM confirmed the presence of 3-layer disks.^31^ It is possible that the 1-layer disk is prone to aggregate into larger species under the heavily perturbation caused by centrifugation, crystallization, or mass spectrometry, making it difficult to observe with these techniques. The cryo-EM data presented here is the first high-resolution structure of a 1- and 3-layer TMVP species. Due to the anisotropic shape of the disks, they all tend to deposit face-down on TEM grids, making them very challenging to distinguish from each other by negative staining. It was previously shown that two different faces can be observed in TEM images, but this was always interpreted as evidence for the 2-layer head-tail disk (AB-disk) rather than disks with an odd number of layers.^32^

**Figure 4.**
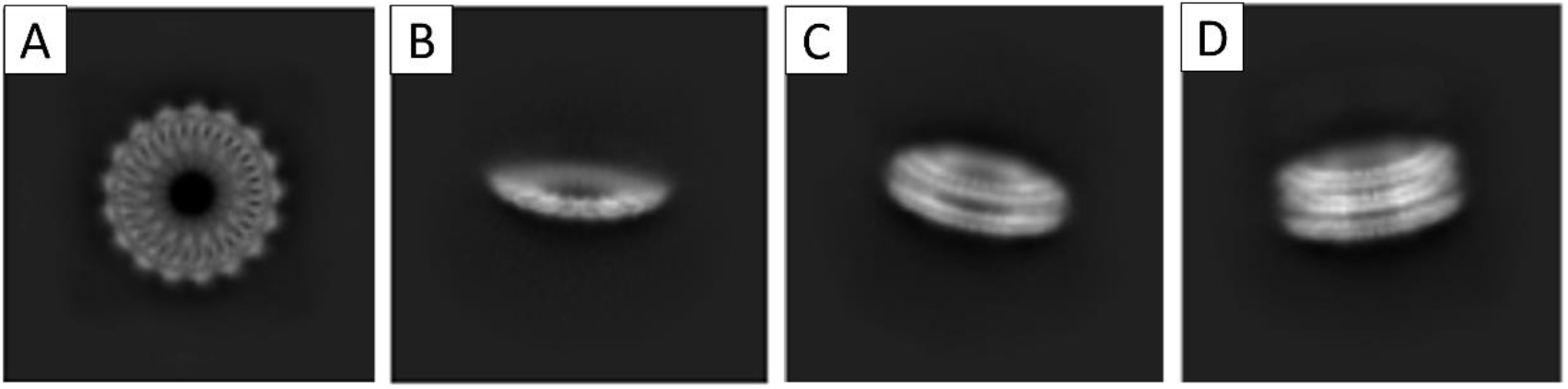
2D class averages of pTMVP disks from cryo-EM. (A) Face view of a disk. (B) Side view of 1-layer disk. (C) Side view of 2-layer disk. (D) Side view of 3-layer disk.

Analytical centrifugation previously found a sedimentation coefficient of 28S that matched the theoretical value for a 3-layer species. However, the authors chose not to make that assignment because the species had never been confirmed in TEM.^10^ The 28S species was instead assigned to a stack of two 2-layer disks, although the theoretical sedimentation coefficient for such a stack is 34-36. The sedimentation coefficient for a 1-layer disk was not previously considered, but species of 12-13S were occasionally observed, which might represent a 1-layer disk. The identity of the 12-13S species was never determined, but it was expected to have approximately 20 monomers, which is close to the 17 monomers of a 1-layer disk. With strong evidence for the existence of 1- and 3-layer disks, we can reassign the sedimentation coefficients from the literature to match very well with the expected values for each disk species, as shown in Table 1.

**Table 1.** Sedimentation constants from previous studies reassigned based on new cryo-EM evidence.

| **Disk Species** | **Theoretical $S_{20,w}$** | **Old Assigned $S_{20,w}$** | **New Assigned $S_{20,w}$** |
| --- | --- | --- | --- |
| 1-layer | Not Calculated | None | 12-13 |
| 2-layer | 19-20.5 | 19 | 19 |
| 3-layer | 27-29 | None | 28 |
| Stack of two 2-layers | 34-36 | 28 | 37 |
| Stack of three 2-layers | 46-49 | 37 | 45 |
| Stack of four 2-layers | 54-59 | 45 | None |

### 1-Layer pTMVP Disks

The resolved portion of the 1-layer pTMVP disk closely resembles the disks from the PDB model (Figure 5). As with the bTMVP disk, the radius of the cryo-EM model is slightly larger and most of the differences from the PDB model are at the poorly resolved inner radius, which is also where most difference between the literature A- and B-disks exist. The inner radius region is less resolved compared to all other models, likely due to the lack of interaction partners. Lateral contacts appear to be very similar to those in the bTMVP disk. The Arg46 side chain interacts with the backbone oxygen of Gly32 instead of forming interlayer contacts as in the other models. The map indicates that residues 90, 91 and 113-116 near the inner loop have different structure than in the PDB disks. The connecting residues could not be modelled. Some solvent exposed side chains, such as Arg134 and Glu50, show multiple conformations in the cryo-EM map.

**Figure 5.**
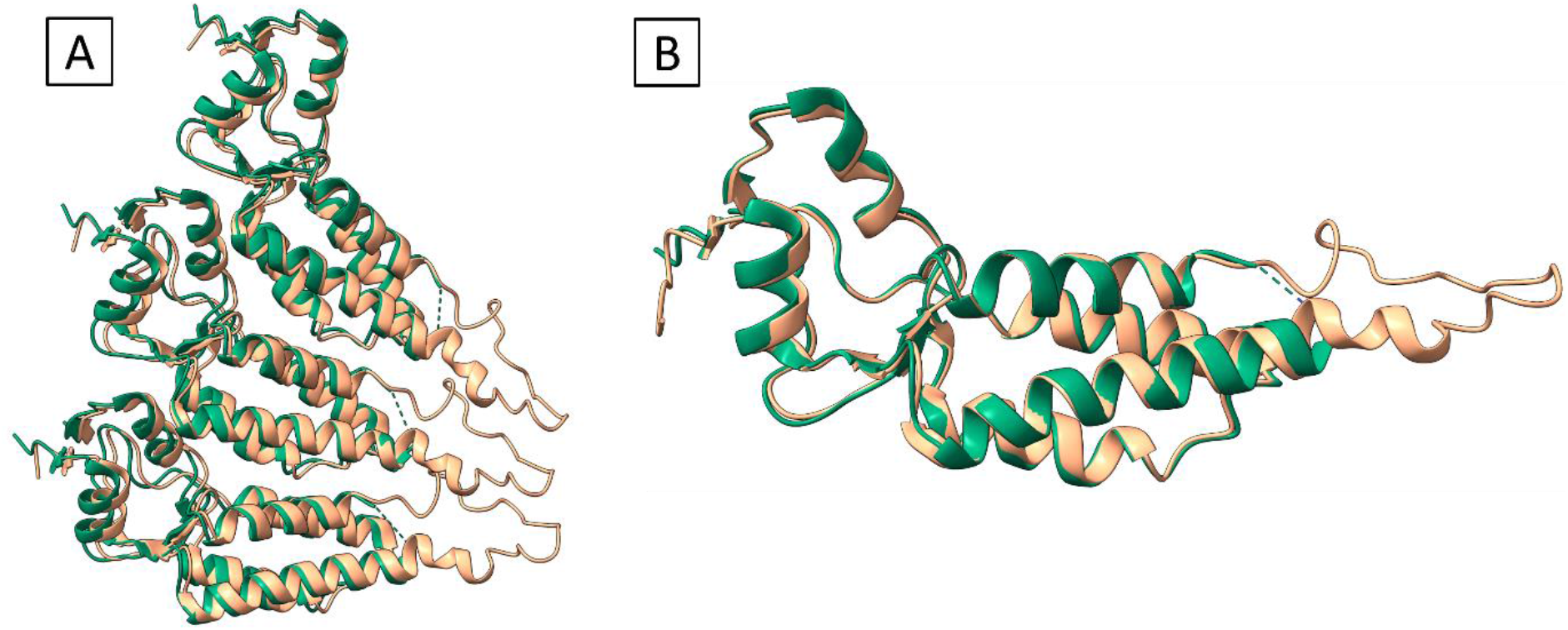
(A) Trimer comparison from aligned pTMVP 1-layer disk and B-disk. (B) Aligned individual monomers showing the divergence at the inner radius. pTMPV coloured green, B-disk coloured peach.

### 2-Layer pTMVP Disks

Despite being a head-head disk, the 2-layer pTMVP disk is not *C*_*2*_-symmetric in solution. The disk has one layer with helices tilted out of the plane of the disk, while the helices in the other layer are flatter (Fig. 6). Due to the unexpected lack of *C*_*2*_-symmetry, this disk may be the same species identified as a head-tail disk in the low resolution cryo-EM study by Butler *et al*. 2D projections of the two disks would appear very similar at low resolution. The backbones of both layers are nearly identical. The main difference in side chain conformations was the presence of a Gln36-Asp115 contact in the convex layer, while Gln36 interacts with Gln38 and Gln34 in the flat layer. Many of the differences between the two layers likely occur in the poorly resolved inner radius region. Unexpectedly, the relative rotation of the two layers of the pTMVP double disk differs by one *C*_*68*_ rotation, or ¼ of a *C*_*17*_, compared to the known AA-disk structure. While this is only a few degrees, such rotations could be relevant to applications that rely on coupling functional components between the layers of the double disk, such as the light-harvesting arrays created by the Francis group.^33^ Presumably the N-terminal acetylation leads to the difference in disk structure and the rotation of the bTMVP disk layers places the C-terminus slightly closer to the N-terminus of the opposing layer, but no direct interactions between the oppositely-charged termini were resolved in the cryo-EM maps.

**Figure 6.**
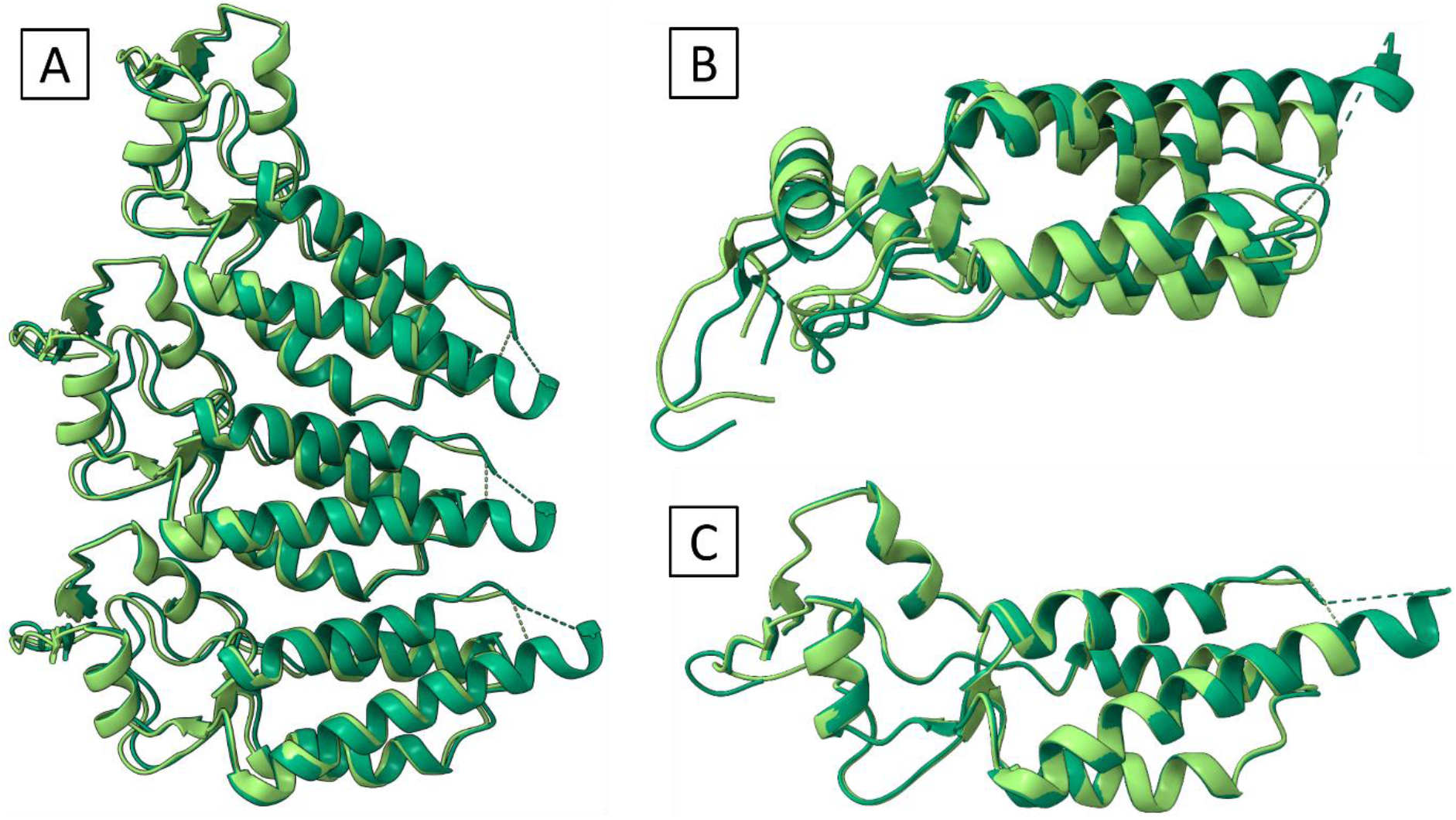
(A) Trimers from aligning the two layers of the pTMVP disk to one another. (B) Side view of monomers from aligned layers showing the difference in quaternary structure between the two layers. (C) Aligned monomers from each layer showing nearly identical secondary/tertiary structure. Flat layer in light green, convex layer in dark green.

The only direct protein-protein contact between the two layers appears to be a Val60-Val60′ interaction near the outer radius. Other interactions between the layers must be solvent mediated, similar to those reported in the 4-layer aggregate. The last three C-terminal residues, 156-158, and the flexible inner loop were poorly resolved, so any interactions at those locations could not be determined. Near the inner radius, the Arg46 side chain is directed towards the opposing layer, but no contacts were observed within 4 Å. The Arg134 side chain on both faces of the disk is bent slightly towards the face and interacts more substantially with Glu131 compared to the bTMVP disk. Due to the difference in rotation of the two layers, Thr59 and Gln57 do not appear to be involved in direct interactions with side chains on the opposing layer unlike what was observed in the bTMVP bilayer disk. Similarly, the Arg112 side chain is aligned closer to the axis of the LR-helix and interacts with Asp115 on the same α-helices. The fact that more protein-protein interactions are observed in the bTMVP disk than in the bilayer pTMVP disk explains why the bTMVP disk is more stable and less prone to making helical rods.

### 3-Layer pTMVP Disks

The layers of the 3-layer disk will be denoted as top, middle, and bottom, as in Figure 7A. The top layer has the opposite orientation of the other two layers. The top and middle layers interact in a head-head fashion, while the middle and bottom layers have a tail-head interaction. The top layer has a backbone and quaternary structure between the A- and B-disk disks. The α-helices bundles in the middle and bottom layers are significantly tilted relative to the plane of the disk compared to the A- and B-disks (Figure 7B). This tilt closely resembles the tilt of the alpha helices in the helical rod model. Lateral contacts near the outer radius are very similar to those in the top layer, but some notable differences are observed near the inner radius. While the 3-layer disk is not helical, the middle and bottom layers appear poised to transition to a helical structure. A 3-layer structure was previously proposed as a possible nucleating species for the helical rod.^30^ Butler observed a 2-turn proto-helix in an early cryo-EM study of TMVP disks at pH 6.8. It seems likely that the 3-layer disk observed here is a precursor to such a proto-helix. Comparing a monomer from the middle layer with monomers from the helical rod, A-, and B-disks reveals that the backbone closely matches the A-monomer with some differences in the inner loop and changes in side chain conformations.

**Figure 7.**
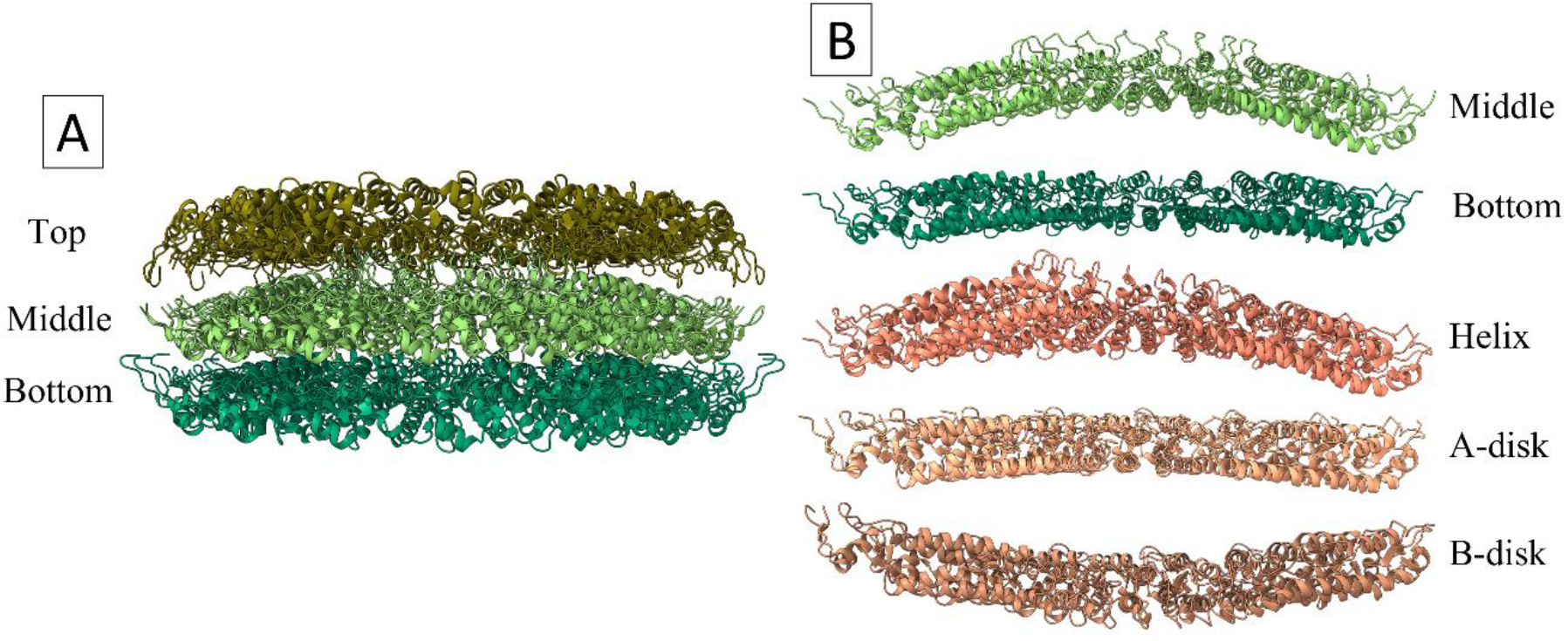
(A) Model of the 3-layer disk with each layer named. (B) Comparison of half-disk models of the middle and bottom layers with selected literature models.

Interactions between the top and middle layers are primarily between residues of the RS-helix in each layer (Fig. 8). In the top layer, Arg46 and Gln47 are involved in numerous interactions with residues in the middle layer. Gln39 in the top layer interacts with Gln39′ and Gln38′ in the middle layer. Asn29 and Gln47 in the top layer interact with Arg46′ in the middle layer. Glu97 in the middle layer inner loop might form a contact with Arg90 in the top layer, although the position of the Glu97 side chain is not well-resolved. There appears to be a hydrophobic contact between Val43 side chains in the top and middle layers. Although the inner loop of the top layer was not resolved, it must adopt a structure closer to the A- or B-disk to avoid occupying the same space as the middle layer inner loops. Extensive protein-protein contacts between the middle and bottom layers exist from the outer to middle radius. At the outer radius, residues in the loop near the N-terminus of the middle layer interact with the loop connecting the RS- and RR-helices in the bottom layer. The RS-helix of the bottom layer interacts with the LR-helix of the middle layer. Arg46′′ in the bottom layer and Arg134′ from the middle layer are involved in many contacts between the two layers. Gln47′′ in the bottom layer also appears to be central to many interactions. A more exhaustive description of interlayer contacts can be found in the SI.

**Figure 8.**
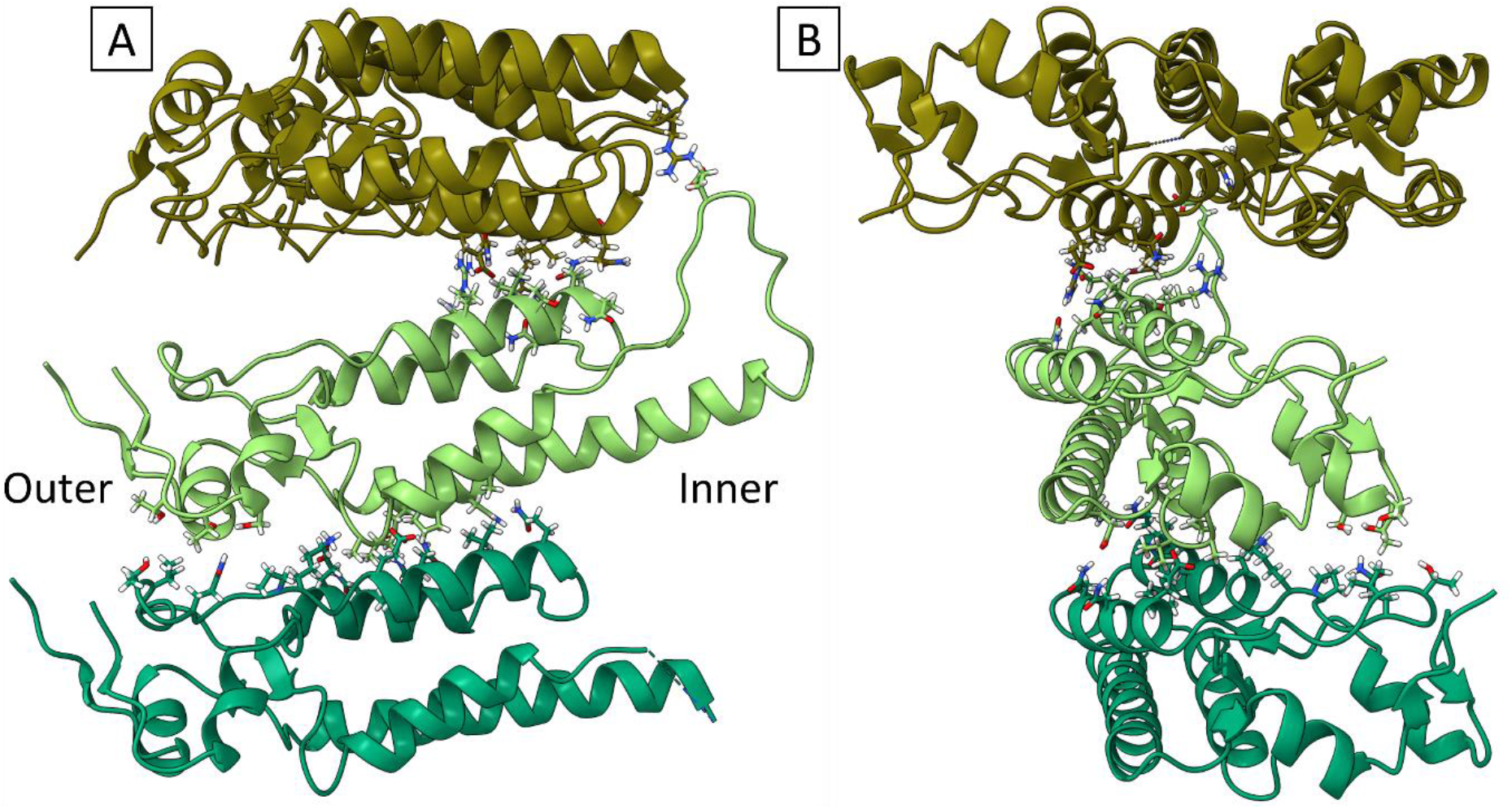
Axial protein-protein contacts of the 3-layer pTMVP disk. (A) Side view of 3-layer interfaces. (C) Side view of 3-layer interfaces. Each layer is coloured a different shade of green.

The positions of the Caspar carboxylates present between the middle and bottom layers were examined. The Glu50-Asp77 pair appear to adopt a similar conformation as reported in the native virus.^14,34^ However, the surrounding residues show different conformations, particularly Lys53′′, Arg46′′, and Thr81′, which seem to interact more with the carboxylate pair in this structure than in the virus. Asp115, which has been found to interact with Arg113 and RNA in the virus, interacts with Arg113′ in the middle layer and Thr89′′ in the bottom layer. In the middle layer, Glu95′, Glu106′, and Asp109′ all form salt bridges with different arginine side chains. As previously discussed, Glu97′ likely forms a salt bridge with Arg90 in the top layer at the head-head interface. In the native virus, only Glu95 was reported to form a salt bridge with an arginine, but it was Arg112 in the adjacent turn of the helix, while here it interacts with Arg90′ on the same subunit. Asp109′ and Asp116′ sandwich Arg112′ in a salt bridge. In the bottom layer, a similar salt bridge between Asp109′′, Asp116′′, and Arg112′′ was observed. Arg113′′ also interacts with Asp116′′. Similar carboxylate-arginine salt bridges were observed in the 4-layer crystal structure.^13^

The resemblance of the middle and bottom layers of the 3-layer disk to the helical rod suggests that the 3-layer disk may be the species that nucleates the proto-helix upon interaction with viral RNA or a reduction in pH. The 1- and 2-layer disks could provide a reservoir for formation of the 3-layer disk and be involved in growth of the helical rod. The head-head interaction between the top and middle layers may serve to stabilize the near-helical conformation of the middle and bottom layers. Butler *et al*. reported that viral RNA is protected in quantized segments corresponding to one or two turns of the helix during virus assembly, supporting the addition of 1-layer disks during the growth of the helical rod.^35^ A general schematic of the transition from disk to helix is presented in Fig. 9.

**Figure 9.**
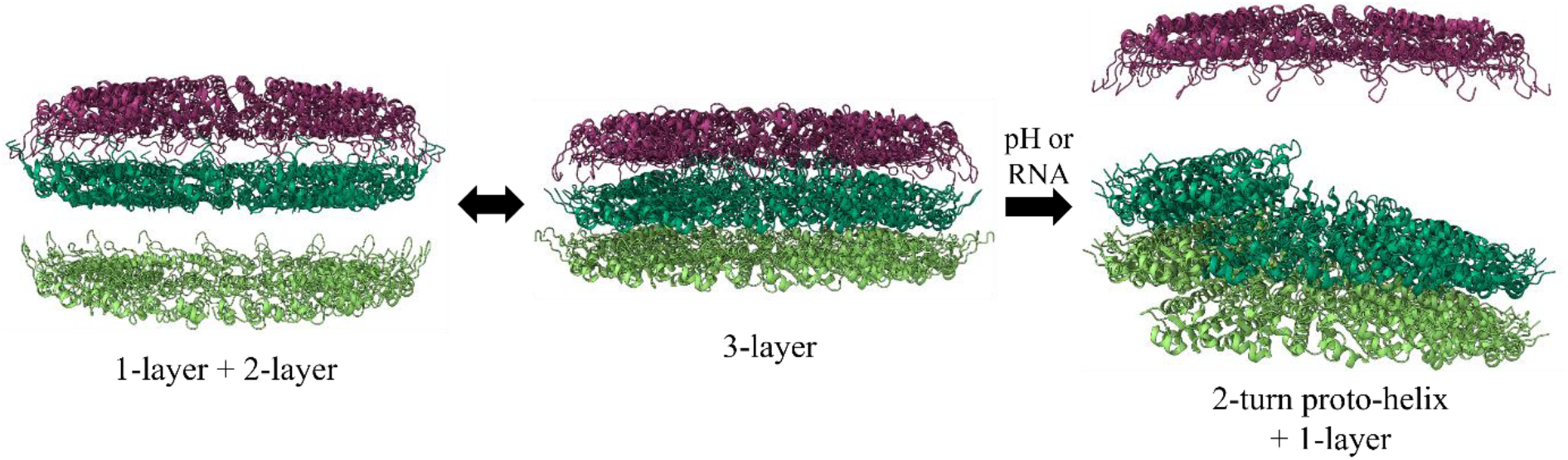
Schematic showing the proposed role of 1-, 2-, and 3-layer disks in the transition to a metastable 2-turn helix, called a protohelix, which nucleates the full helical rod.

## Conclusions

In this work we presented the first high-resolution cryo-EM structures of tobacco mosaic virus disks under conditions known to be favourable for *in vitro* assembly of the virus. TMVP produced in bacteria self-assembled into stacked disk rods after being stored at pH 5.0, which remained intact upon dialysis to pH 7.2. The disks in these stacks closely resemble the AA-disk derived from the 4-layer disk aggregate previously reported in the literature. An interaction between Arg134 side chains on neighbouring disks was well-resolved. An Arg112-Arg112′ interaction at the inner radius is also likely, but not as well-resolved as the Arg134 interaction. Both of these Arg-Arg interactions appear to be stabilized by neighbouring carboxylate residues. No other direct protein-protein interactions were observed between neighbouring bilayer disks in the stack. When bTMVP was dialyzed to pH 7.2 from pH 8.0, a mixture of 1-, 2-, 3-, and 4-layer disks formed. High resolution data is still being collected to allow the reconstruction of detailed models for each of these disk structures. The lack of N-terminal acetylation in bTMVP does not appear to cause any significant changes in secondary/tertiary structure near the termini compared to pTMVP. Additionally, no interactions between the N- and C-termini were resolved, suggesting that the effect of N-terminal acetylation may not be as straight forward as previously thought.

Protein produced in plants showed 1-, 2-, and 3-layer disks. This is the first report of 1-layer disks from wild-type TMVP. The fact that primarily 2-layer disks have been observed in the past likely indicates that 1- and 3-layer disks are less stable and prone to form 2- or 4-layer species during methods such as analytical centrifugation or crystallization. There is also an inherent bias in traditional negative stain TEM, due to the anisotropic shape of the disks, which favours face-down rather than on-edge deposition onto the grid. Disk species observed in cryo-EM match well with sedimentation constants from the literature, although the sedimentation constants were not assigned to 1- or 3-layer disks at the time due to a lack of TEM evidence.^10^ These results also match well with a recent mass spectrometry study of bTMVP disks.^31^ The 2-layer pTMVP disk is not *C*_*2*_-symmetric despite the layers having opposing orientations to each other. The α-helices bundle in one layer is significantly tilted out of the plane of the disk compared to the other layer. It was also found that the relative rotation between the two layers differs from that of the 4-layer crystal structure and the bTMVP stacked disks. The bilayer disk appears to always adopt a head-head structure, but only becomes *C*_*2*_-symmetric when disks are stacked/crystallized, although there is evidence suggesting that a K53R mutation may enforce a head-tail bilayer disk.^36^ There appear to be more solvent-mediated interactions between the layers in the pTMVP bilayer disk than in the bTMVP disk, which may explain the higher stability of the bTVMP disk. The 3-layer disk has two layers with the same orientation and one layer with the opposite orientation. The opposing layer resembles the 1-layer disk, with a structure between the A- and B-disks from the 4-layer aggregate. The α-helices bundle of the middle layer is tilted very significantly, as in the helical rod, but the disk is not helical. The layer with the same orientation as the middle layer shows less severe tilting of the α-helices bundle relative to the plane of the disk. These two layers appear to be poised to transition to a helical structure.

The structure of the 3-layer disk leads us to suggest that the 3-layer disk is the precursor to the proto-helix. The layer with opposite orientation likely stabilizes the two head-tail layers in their near-helical conformation until a reduction in pH or interaction with viral RNA triggers the transition to a stable helical structure. The 1- and 2-layer disks may act as a reservoir for the 3-layer nucleating species in addition to being involved in the growth of the helical rod. The fact that our data agrees well with mass spectrometry results from the Francis group indicates that their charge detection system may provide a more accessible method to characterize the particles present in solutions of TMVP with reasonable accuracy, as cryo-EM is not as widely available as mass spectrometry. The maps and models presented here should facilitate a greater understanding of TMVP behaviour in solution and prove useful for future mutational studies.

## Acknowledgements

We thank Dr. Mike Strauss, Department of Anatomy and Cell Biology, McGill University, for assistance collecting preliminary cryo-electron microscopy data of bTMVP disks.

GC and ASB were supported by the Canada Foundation for Innovation (CFI), Québec Centre for Advanced Materials (QCAM), and Natural Sciences and Engineering Research Council (NSERC).

We gratefully acknowledge Polish high-performance computing infrastructure PLGrid (HPC Center: ACK Cyfronet AGH) for providing computer facilities and support within computational grant no. PLG/2023/016577.

This publication was partially developed under the provision of the Polish Ministry and Higher Education project “Support for research and development with the use of research infra-structure of the National Synchrotron Radiation Centre SOLARIS” under contract no 1/SOL/2021/2.

At the John Innes Centre (JIC) this work was supported by the United Kingdom Biotechnology and Biological Sciences Research Council (BBSRC) Grants BB/T004703/1 and BB/Y005732/1, the Institute Strategic Programme Grant “Harnessing Biosynthesis for Sustainable Food and Health (HBio; grant number BB/X01097X/1), and the John Innes Foundation. We also thank the JIC Horticultural Services department for assistance in the production of plant-derived TMV CP.

## Notes

### Competing Interest Statement

The authors have declared no competing interest.

