## Supplementary Information for "High-Resolution Structures of Tobacco Mosaic Virus Disks from Cryo-Electron Microscopy"

### Materials & Methods

All materials used were reagent grade or better. Sodium phosphate, potassium phosphate, EDTA, and sodium borate were purchased from Fisher Scientific. Terrific Broth and LB media were purchased from MP Biomedicals. Ampicillin and chloramphenicol were purchased from Research Products International (RPI).

### bTMVP Expression

A pET20b vector encoding the sequence of WT-TMVP was purchased from NorClone Biotech by reverting an S123C mutant plasmid kindly gifted by Prof. Matthew Francis (UC Berkeley) back to the wild-type sequence. The 6H-TMVP, 6H-2R-TMVP, and 4H-TMVP plasmids were purchased from NorClone Biotech based on the WT-TMVP plasmid. For all mutants, Tuner(DE3)pLysS competent cells (Novagen) were transformed with the vector and streaked on an LB agar plate supplemented with 100 ug/mL ampicillin and 34 ug/mL chloramphenicol. After overnight incubation at 37 ºC, a single colony was used to inoculate 10 mL of LB and grown overnight with constant shaking at 37 ºC. This saturated growth was used to make frozen glycerol stocks of transformed cells stored at -80 ºC. For a typical expression, 10 mL of LB media supplemented with 100 ug/mL ampicillin and 34 ug/mL chloramphenicol was inoculated with a small portion of cells from the frozen glycerol stock and grown overnight at 30 ºC with constant shaking at 250 RPM. A 1 mL aliquot of the resulting culture was used to inoculate 1 L of Terrific Broth, grown at 37 ºC until $OD_{600} = 0.6$, and then grown overnight at 30 ºC. No isopropylthio-β-galactoside (IPTG) was necessary for high levels of protein expression due to a combination of leaky expression of the promoter and compromised cell growth upon induction by IPTG.[1,2] Cell pellets were harvested by centrifugation and frozen at -80 ºC.

**bTMVP Purification**

Frozen cell pellets were thawed on ice, resuspended in cold lysis buffer (20 mM triethanolamine, 1 mM EDTA, pH 8.0), and lysed by sonication at 50% duty cycle and 60% amplitude for 5 minutes on ice. The resulting lysate was clarified by centrifugation at 32000 RCF for 30 minutes at 4 ºC. The supernatant was collected and allowed to reach room temperature. Solid ammonium sulfate was added slowly with stirring to a final concentration of 35% (v/v). The resulting precipitate was isolated by centrifugation at 32000 RCF for 30 minutes 4 ºC and resuspended in minimal lysis buffer. The solution was then dialyzed overnight at 4 ºC against 4 L lysis buffer to remove residual ammonium sulfate. Any remaining precipitate was removed by centrifugation. The solution was diluted to a final volume of 250 mL with lysis buffer and loaded on a DEAE Sepharose anion exchange column (Cytiva) overnight at 4 ºC. The protein was eluted with a 0-300 mM NaCl gradient using an AKTA Avant FPLC. Fractions showing absorbance at 280 nm were analyzed by Tris-buffered SDS-PAGE on an 8-16% gradient gel (Biorad) run in constant voltage mode at 200 V and stained with Coomassie Blue R250 (Fig. S10). Pure fractions were combined and concentrated. Pure protein was dialyzed at 4 ºC against 20 mM sodium borate buffer at pH 8.5, concentrated to 2.7 mg/mL, and filter sterilized. Aliquots were frozen in LN2 and stored at -80 ºC until further use.

**pTMVP Expression and Purification**

TMV nanorods were purified *Nicotiana benthamiana* leaves infiltrated with construct pEff-TMV-CP/OAS.[3,4] The coat protein was isolated using the acetic acid method and quantified using uv/vis spectroscopy at A280.[5,6] The coat protein was stored in the form of helices at pH 5.0 prior to dialysis to the required pH for cryo-EM analysis.

**Cryogenic Electron Microscopy**

**pTMVP Disks**

4 uL of a solution of TMVP at 2.5 mg/mL in 75 mM sodium potassium phosphate pH 7.2 was applied to a glow discharged grid and plunge frozen in liquid ethane using an FEI Vitrobot. 8152 micrographs were collected using a Titan Krios cryo-microscope with a Falcon III camera at 75k magnification. All micrographs were motion-corrected using MotionCorr2 and CTF estimation was performed using PatchCTF in cryoSPARC v4.3.0.[7] Manual particle picking was performed on 100 micrographs. Approximately 250 particles were extracted and classified into 2D classes. These classes were then used for another round of particle picking and 2D classification using template picker from 500 micrographs. The best classes were used as a template for template picker on the full set of micrographs. 2D classes from the full data set were used to train a Topaz neural network. Two rounds of training were performed. Each training was followed by particle extraction, and 2D classifications to select the best-looking classes. *Ab-initio* reconstruction was performed on 231,671 particles from 5 classes. The best 3 classes were subjected to homogeneous refinement with C1 symmetry followed by another round of homogeneous refinement with $C_{17}$ symmetry. Final maps had resolutions of ~3 Å based on FSC at 0.143 threshold.

**bTVMP Stacked Disks**

4 uL of a solution of TMVP at 2.5 mg/mL in 75 mM sodium potassium phosphate pH 7.2 was applied to a glow discharged grid and plunge frozen in liquid ethane using an FEI Vitrobot. 8488 micrographs were collected using a Titan Krios cryo-microscope with a Falcon III camera at 75k magnification. All micrographs were motion-corrected using MotionCorr2 and CTF estimation was performed using PatchCTF in cryoSPARC v4.3.0.[7] Approximately 300 particles were manually picked and then extracted and classified into 2D classes. These classes were then used for another round of particle picking and 2D classification using the filament tracer tool from 500 micrographs. The best classes were used as a template for filament tracer on the full set of micrographs. Two rounds of particle picking were performed on the full dataset using the best classes from the previous round. Helical reconstruction was performed on 131,946 particles, yielding a map with a resolution of ~2.87 Å based on FSC at 0.143 threshold. Helical refinement was performed on 68,578 particles to give the final map with a resolution of 2.24 Å based on FSC at 0.143 threshold.

**Model Refinement**

Refinement began using a monomer from the B-layer of PDB 1EI7 with or without an N-terminal acetyl group. Rigid-body fitting was performed in ChimeraX.[8] The models were refined through iterative rounds of automated refinement with Phenix real.space.refine and manual adjustment using the ISOLDE tool in ChimeraX.[9,10] All figures were created in ChimeraX.

**Figures**

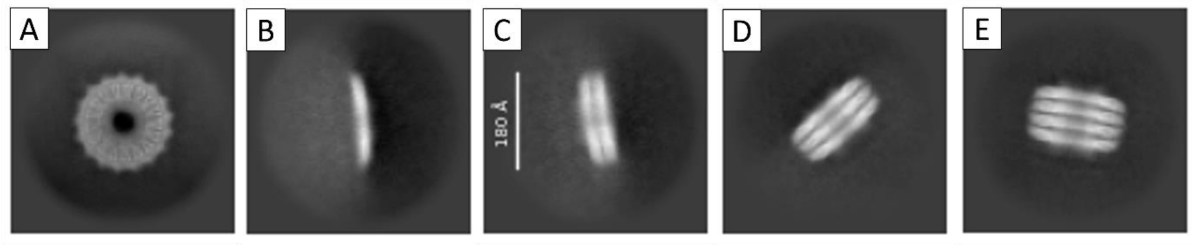


**Figure S1:** Preliminary 2D class averages of bTMVP disks stored under alkaline conditions. (A) Face view. (B) 1-layer side view. (C) 2-layer side view. (D) 3-layer side view. (E) 4-layer side view.

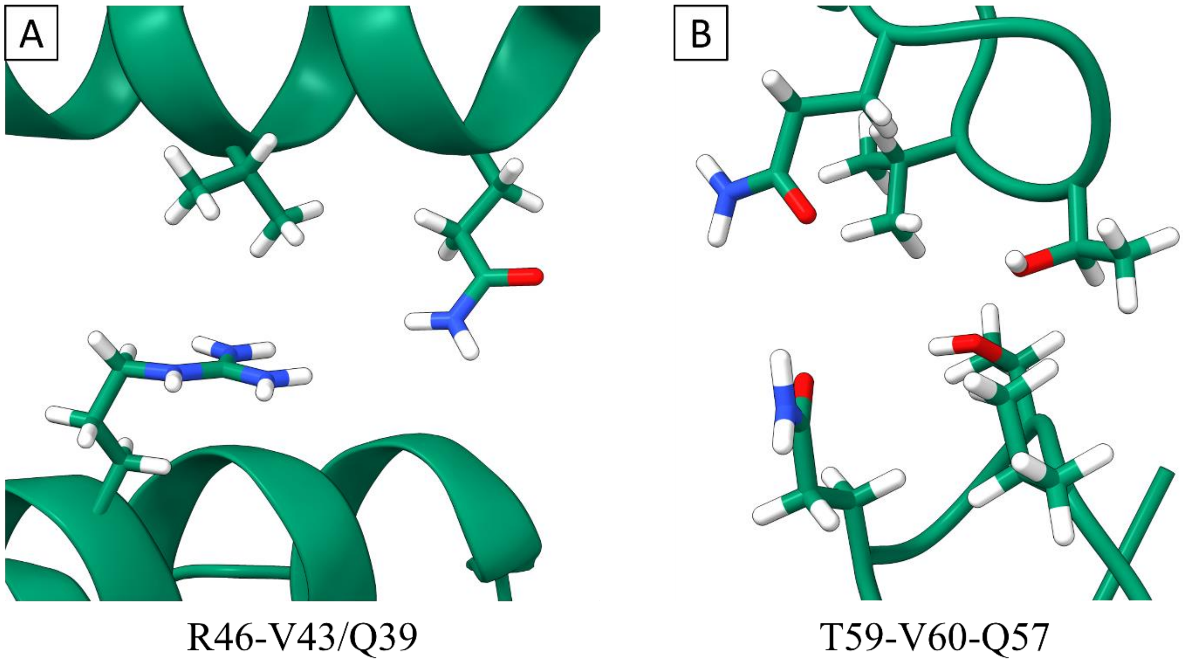


**Figure S2.** Head-head contacts in the bTMVP bilayer disk. (A) R46-V43′/Q39′. (B) T59/V60/Q57-T59′/V60′/Q57′.

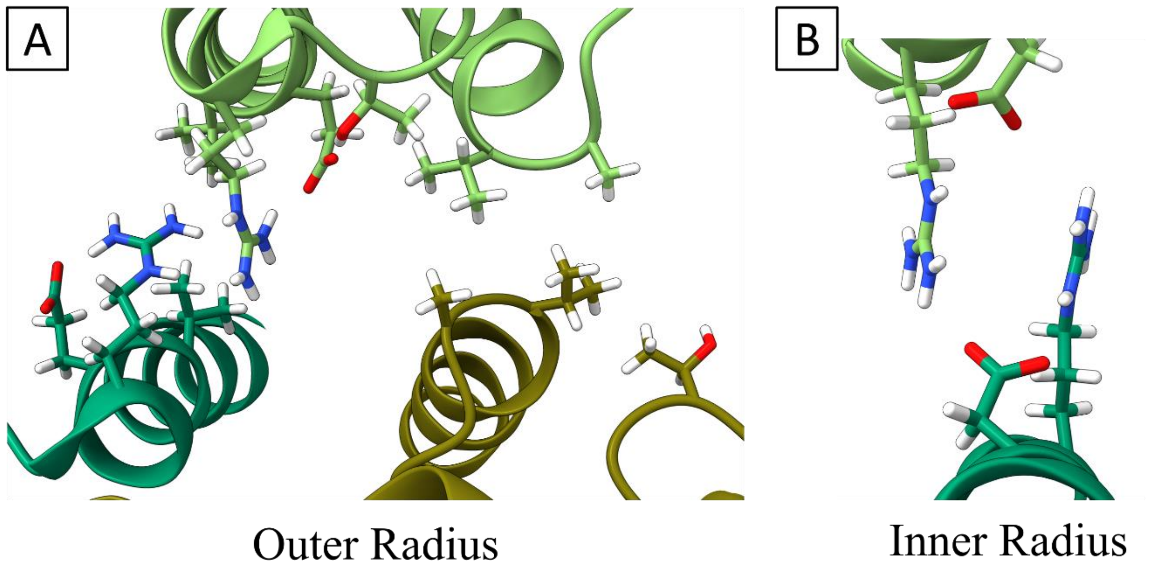


**Figure S3.** Tail-tail contacts in the bTMVP stacked disks. (A) Outer radius clusters of V130/E131/R134-V130′/E131′/R134′ and V75/A74-A74′/V75′ with T136 bridging between the clusters. (B) Inner radius contacts of R112/D116-R112′/D116′.

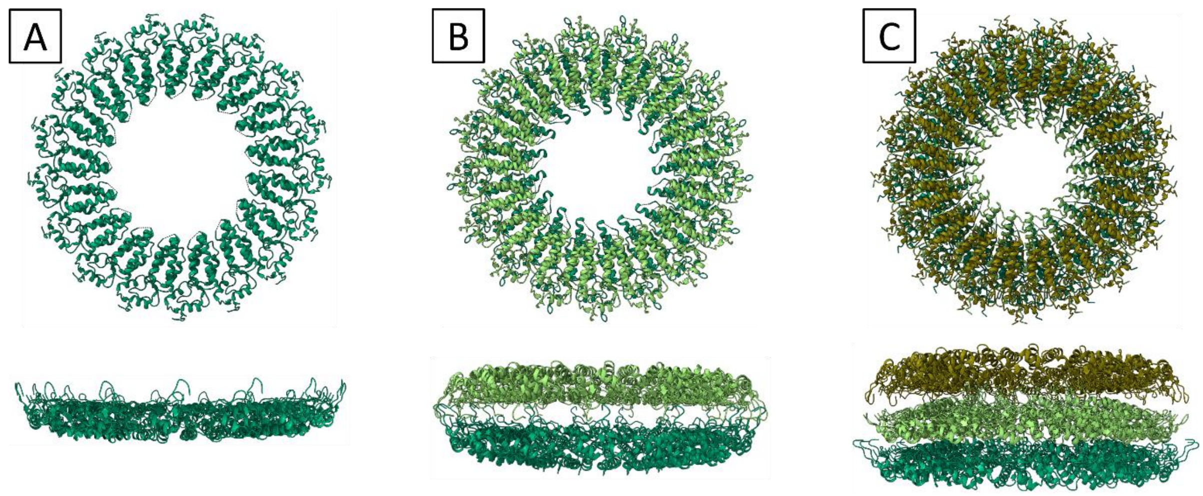


**Figure S4.** Face and side views of refined pTMVP disk models. (A) 1-layer. (B) 2-layer. (C) 3-layer.

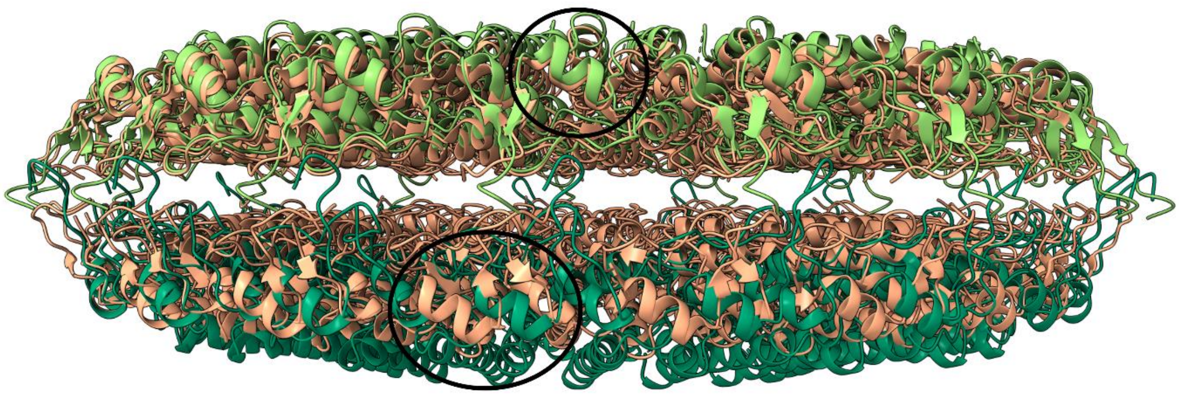

**Figure S5.** Comparison of the relative rotation in the layers of pTMVP (green) and AA-disk (peach). Black circles indicate matching α-helices in top layers and mismatched α-helices in bottom layers.

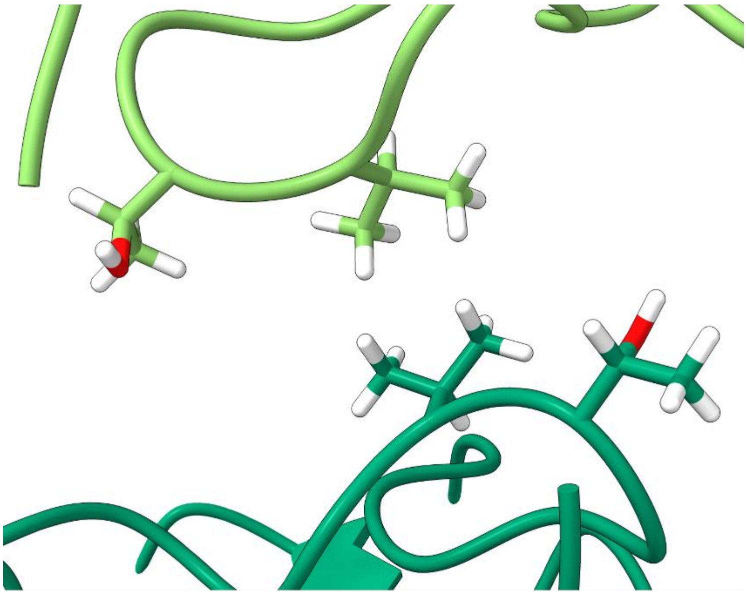

V60-V60

**Figure S6.** V60-V60′ contact pair at the outer radius of pTMVP 2-layer disk.

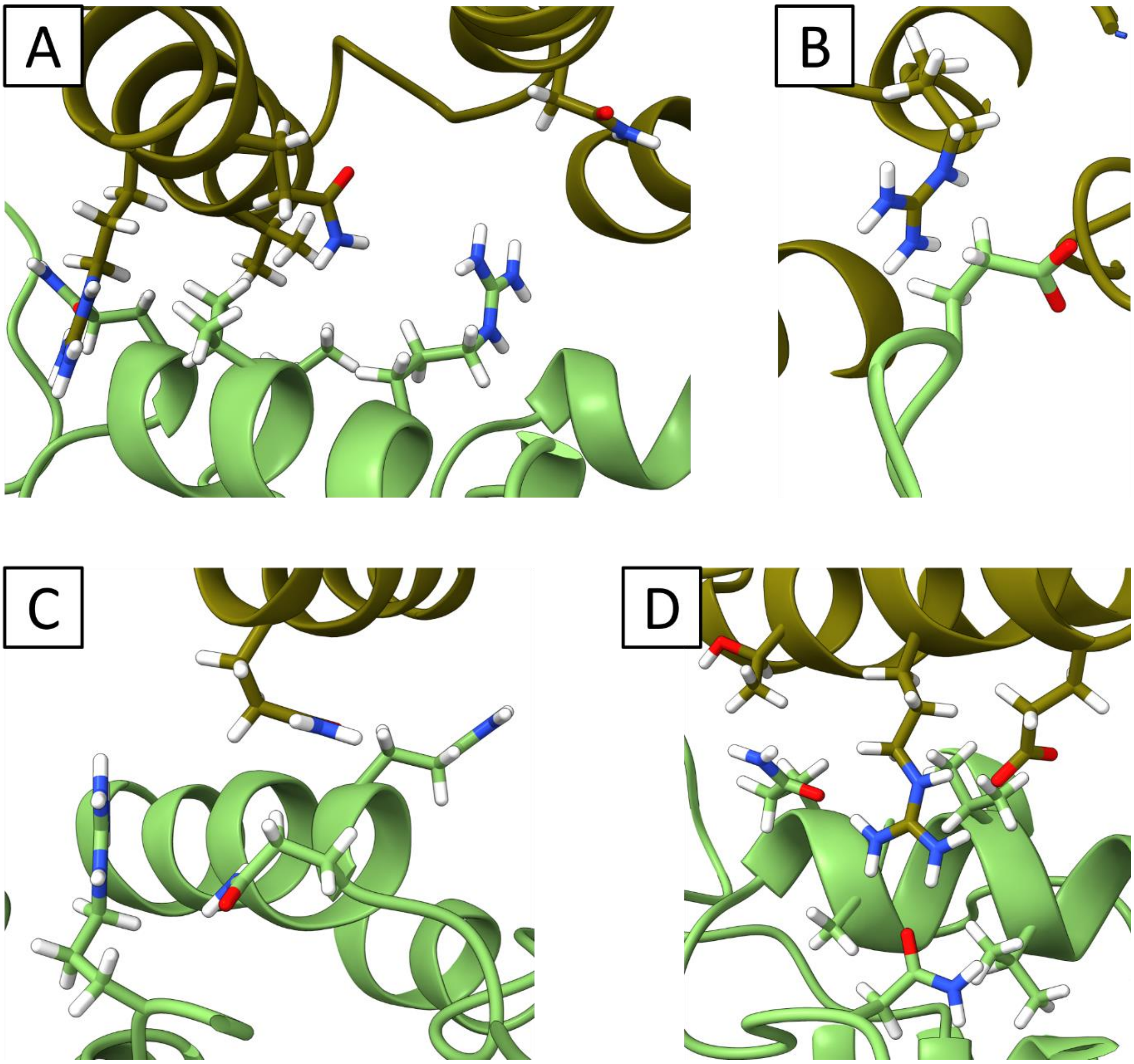


**Figure S7.** Contacts between the top and middle layers of 3-layer pTMVP disk.

(A) Q47/N29/V43/R46-R46′/V43′/T42′/Q39′. (B) R90-Q97′. (C) N39-N39′/N39′/R90′.

(D) R46/E50/T42-N33′/Q39′/A40′/V43′/V44′.

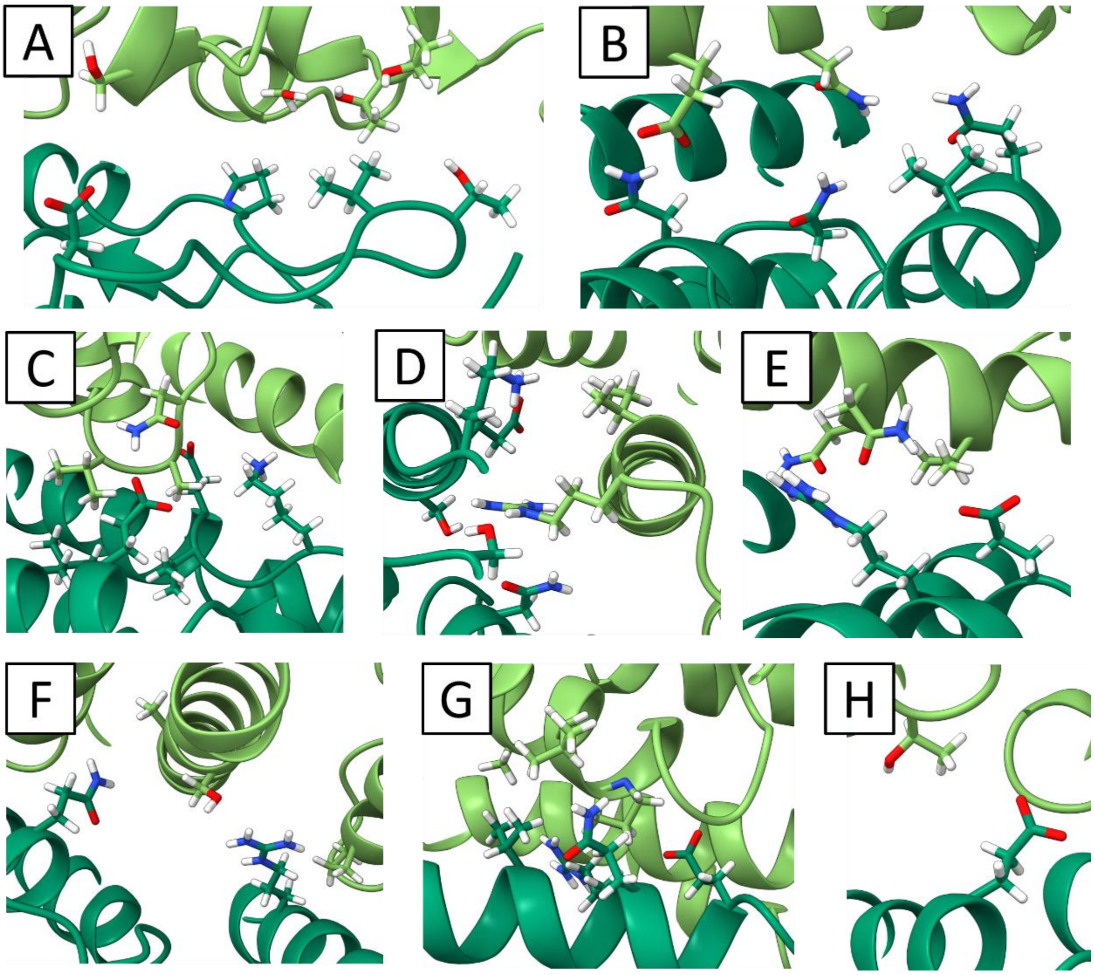


**Figure S8.** Contacts between the middle and bottom layers of 3-layer pTMVP disk. (A) T5/T6/S8/S146-P54′/Q57′/T59′/V60′/D66′. (B) N127/E131-N29′/N33′/Q39′/V43′. (C) N73/A74/V75-E22′/L26′/E50′/V51′/K53′. (D) I133/R134-S15′/N29′/S49′/E50′/K53′. (E) N126/V130-R46′/E50′. (F) P78/S123/A124-Q39′/R46′. (G) P78/L79/A82-Q39′/V43′/R46′/Q47′/E50′. (H) T136-E22′.

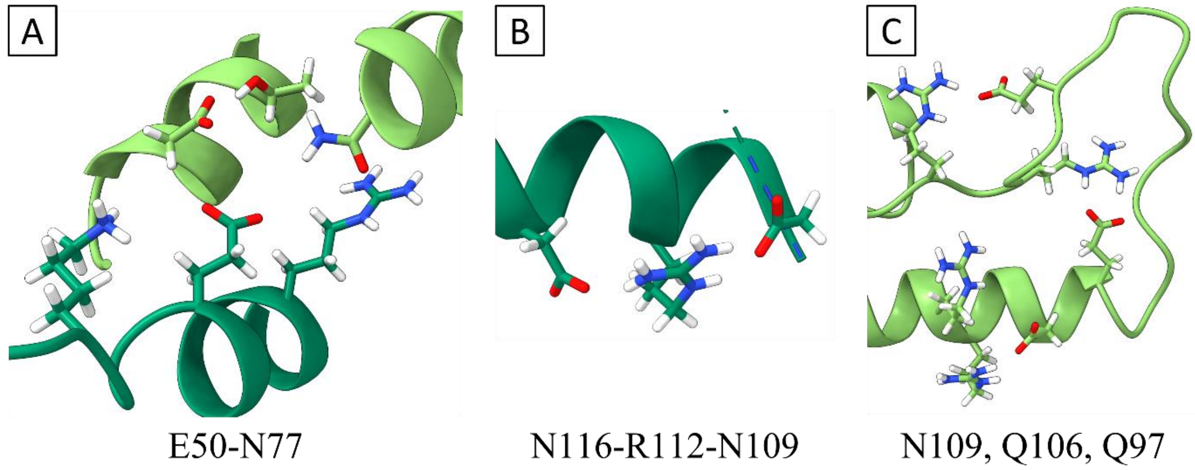


**Figure S9.** pTMVP 3-layer disk Caspar carboxylates. (A) E50-N77′ with neighbouring K53, R46, T81′, and N126′. (B) E109-R112-E116 in the bottom layer. (C) Carboxylate-arginine salt bridges at the inner radius of the middle layer.

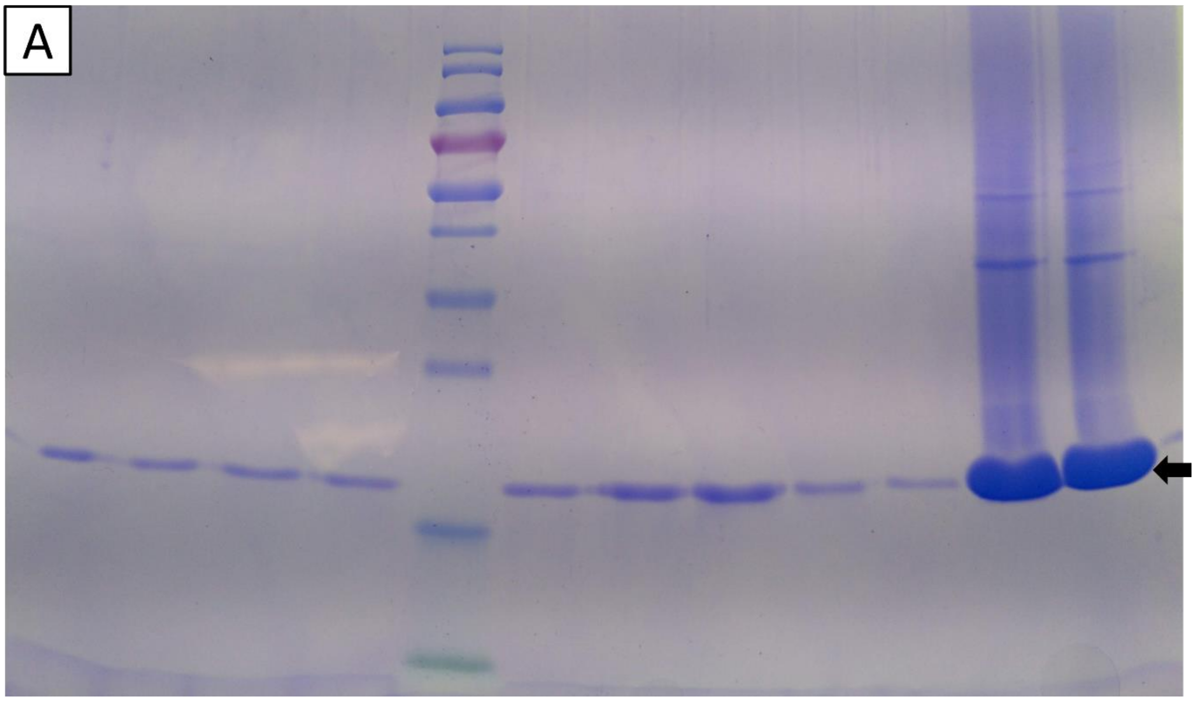


**Figure S10.** SDS-PAGE of TMVP column elution. Molecular weight marker: 10 (green), 15, 25, 35, 40, 55, 70 (red), 100, 130, 180 kDa. Arrow indicates target protein band. Higher molecular weight bands appear to correspond to dimer and trimer molecular weights.
